# Precise Memory is Predicted by Learning-Induced Sensory System Neurophysiological Plasticity

**DOI:** 10.1101/600866

**Authors:** Elena K. Rotondo, Kasia. M. Bieszczad

## Abstract

Despite identical learning experiences, individuals differ in the memory formed of those experiences. Memory formed with sensory specificity determines its utility for selectively cueing subsequent behavior, even in novel situations. If an individual forms generalized memory, then there is potential for novel sensory cues to interfere with accurate behavioral performance. Here, a rodent model of auditory learning capitalized on individual differences in learning-induced auditory neuroplasticity to identify and characterize neural substrates for sound-specific (vs. general) memory of the training signal’s acoustic frequency. Animals with naturally or pharmacologically induced signal-“specific” memory revealed behaviorally, exhibited long-lasting signal-specific neurophysiological plasticity in auditory cortical and subcortical evoked responses, while learning-induced changes were not detected in animals with “general” memories. Individual differences validated this brain-behavior relationship, such that the degree of change in neurophysiological responses could be used to determine the precision of memory formation.

## Main Text

Individual differences are often intentionally minimized by experimental designs in behavioral neuroscience. Consequently, traditional approaches implicitly treat differences between subjects as “noise” to be overcome by large enough sample sizes to accurately describe and compare the underlying distributions of each group. This philosophical choice stymies an opportunity to use variability as a key to understand brain-behavior relationships. Even when group analyses reveal significant effects, they may not identify the full extent of brain-behavior relationships. Variability in learning, memory, and behavior derives from multiple factors^1-8^ each with their own neural generators that have distinct effects on function^9-11^. Therefore, a powerful use of within-group variability is to explain magnitude-of-effect in individual behavioral performance by differences in learning-induced neural function. Indeed, a major goal in behavioral neuroscience is to understand brain-behavior relationships enough to gain the ability to manipulate function and drive behavior in a desired direction by promoting neuroplasticity^12-14^. However, to harness plasticity mechanisms in a useful way after the form of experience-induced plasticity between groups is identified, its function and magnitude of effect must be determined at the level of the individual.

Previous work has identified distinct functions for different forms of auditory system plasticity related to the behavioral relevance of acoustic frequency^6,15-18^ including at different time scales^19-21^. For example, in the auditory cortex, rapid changes in neural tuning properties correlate with selective attention to sound frequency during active tasks^21,22^, while long-term changes in tuning bandwidth may be related to frequency-specific memory over time^23,24^. Subcortical neurons in the lemniscal auditory nuclei also exhibit changes in tuning properties to represent behaviorally relevant sound frequencies across the lifespan^25,26^. Furthermore, the profound auditory cortical modulation of subcortical sound processing^25,27,28^ suggests that auditory system plasticity interacts at multiple levels to serve high-order functions beyond simple feature coding of acoustic frequency. As such, non-invasive auditory brainstem neurophysiological responses (ABR) in humans that captures system-wide processing of sound can predict individual high-order auditory skills that involve listening-in-noise and language intelligibility^29,30^.

Importantly, adult learning-induced cortical plasticity is not merely a reflection of encoded sound-stimulus statistics; instead it reflects task-related rules that support adaptive behavior^30,31^. Furthermore, and perhaps the strongest reason for extending research to the individual, is that individual subjects seldom form identical memory, even within a group that has undergone the same sound-frequency training with *identical* task rules^5,12,33^.Therefore, other factors must explain variable outcomes. However, such individual differences create an opportunity to discover neural plasticity that can account for behavioral functions. For example, a neural change in individual subjects that matches the specific sensory content of the individual subject’s memory can validate the candidate as a neural substrate of those specific contents. Thus, the auditory system may provide a substrate for the specificity of acoustic content of memory, albeit only one part of the complete memory, which is likely distributed in other modalities of the brain and in various forms.

Here, the auditory content of memory is defined in the highly-tractable acoustic frequency domain, where animals may remember a sound cue *specifically* as the particular frequency heard during training with reward. A precise memory for acoustic frequency is *specific* to the acoustic frequency of the training signal. This is in contrast to memory for sound that is *generalized* across acoustic frequency, in which animals do not remember acoustic frequency *per se*, as the mere presence of a sound signal will effectively cue behavior. Given that learning experiences with sound can alter the frequency-tuning properties of both cortical and subcortical auditory neurons, we set out to determine whether learning-induced auditory system plasticity in cortical and subcortical areas are neural substrates for the frequency-specificity of memory. We predicted that a form of frequency-specific neurophysiological plasticity could emerge at a group-level analysis, while individual differences would validate its behavioral function for frequency-specific memory.

## Results

To determine the function of auditory system plasticity for memory, auditory brainstem responses (ABRs) were recorded from a group of 6 adult, male rats before and after they learned a simple single-tone auditory operant task (tone-reward training; see Methods). Rats were trained with a 5.0 kHz signal tone that predicted availability of a water reward after an operant response (bar-press) (Fig. 1a). After reaching asymptotic performance, animals were given a memory test (Fig. 1b). In this test, rats were randomly presented with the 5.0 kHz signal tone and four novel tones that differed in acoustic frequency to determine which individuals responded selectively only to the trained sound frequency (specific memory) vs. which individuals that responded broadly across acoustic frequency (general memory). Despite being trained identically in this single-tone task, animals varied in the distribution of responses among memory test tone frequencies. To identify a potential candidate form of early auditory system plasticity for memory, the change in response amplitude of the first peak (PW1) in the signal tone-evoked ABR was calculated for each individual from a recording session outside of the training context, which was made once before training, and again after reaching high levels of performance in training (Fig. 1c). One animal (n=1/6) that responded selectively only to the trained sound at Memory Test, is said to have remembered the signal tone (5.0 kHz) *specifically*. This animal showed the largest PW1 peak amplitude increase (M=71.64%), while other animals showed less specificity (n=5/6) and variably smaller changes in PW1 amplitude (M=-1.01, SE=13.52). Importantly, comparing individual PW1 amplitude changes against an animal’s own proportion of bar presses made to the signal tone (vs. novel tones) during the memory test revealed a significant positive correlation: the greater the increase in peak amplitude, the greater the specificity of behavioral responses to the signal tone at memory test (*r=0.842, p=0.035*). These findings support a function of this form of ABR plasticity is to underlie long-term sound-specific memory. A subsequent experiment capitalized on a known molecular mechanism of long-term memory formation^34,35^ that we hypothesized would drive interacting forms of plasticity across the auditory system towards memory substrates for more frequency specificity and selective behavior.

**Fig. 1.**
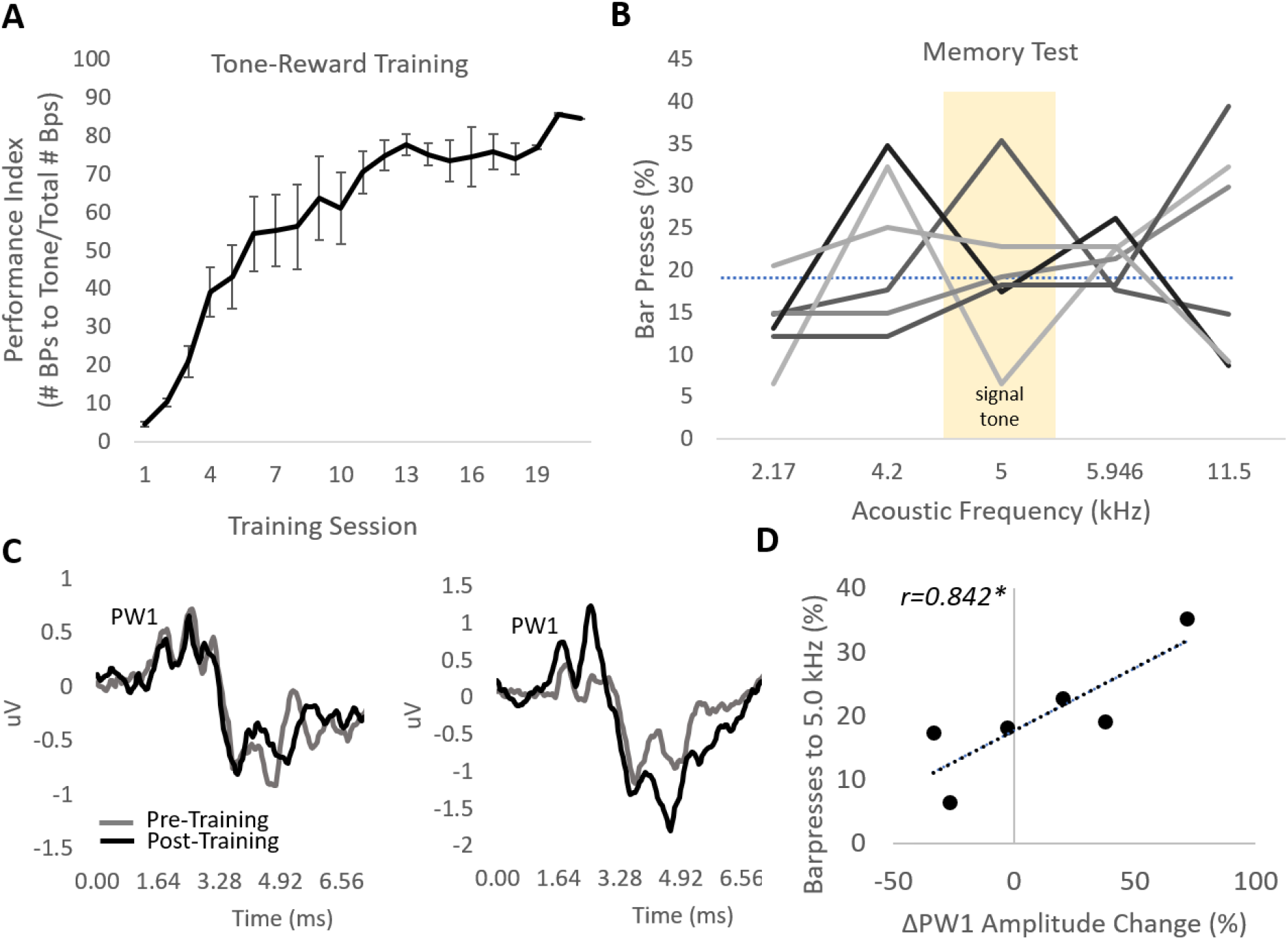
Individual differences in auditory brainstem response plasticity predict sound-specific memory following identical learning experience. (A) Rats learn to associate a 5.0 kHz signal tone with an operant water reward. Individuals are trained to asymptotic performance. Error bars represent +/- SEM. (B) Despite being trained with a single tone, the Memory Test reveals variability in how selective responses are to the training tone frequency (5.0 kHz). In fact, only one rat responded substantially more the signal tone vs. other tone frequencies. Lines represent individual subjects. (C) Tone-evoked auditory brainstem responses recordings were obtained before (“pre-training,” gray) and after (“post-training,” black) to determine learning induced changes in signal tone-evoked positive wave 1 (PW1) amplitude. Graphs depict data from 2 of 6 individuals. (D) There is a significant positive correlation between individual differences in PW1 amplitude changes and the percent of responses made to the signal tone during the Memory Test. Greater amplitude increases predict a greater percent of responses allocated to the signal tone frequency. *p<0.05

Epigenetic mechanisms, such as histone deacetylase 3 (HDAC3), have been recently studied for their strong influence on the formation, strength, and persistence of long-term memory^36^, including for sensory cues^37^. Inhibition of class I HDACs like HDAC3, which releases HDAC-mediated constraints on gene expression, has been found to enhance the specificity of auditory memory across species^24,34,38^ including humans^39^. Here, we confirm that HDAC3 inhibition enhances the frequency-specificity of auditory memory (Fig. 2). Rats given injections of the HDAC3 inhibitor, RGFP966 (Abcam Inc., 10 mg/kg, n=6; vs. vehicle, n=7) immediately following early sessions of tone-reward training exhibited a frequency-specific behavioral response distribution at Memory Test weeks later (Fig. 2a), which characterizes a “specific” memory phenotype. RGFP966-treated animals responded significantly more to the 5.0 kHz signal frequency tone than to distant tones (M=20.04, SE=6.67, one-sample t-test: t(5)=3.00, p=0.029), while the difference did not reach significance to nearby tones (M=19.26, SE=9.65, one-sample t-test: t(5)=1.99, p=0.102) (Fig.2b). In contrast, vehicle-treated animals did not behaviorally discriminate the signal tone from nearby tones (M=0.94, SE=7.14, one-sample t-test: t(6)=0.132, p=0.899), nor from distant tones (M=-4.82, SE=8.45, one-sample t-test: t(6)=-0.5711, p=0.588), which characterizes a “general” memory phenotype. Compared to untreated rats (n=1 out of 6 individuals), RGFP966 treatment significantly increased the proportion of individuals in the group with “specific” memory (to n=4 out of 6, binomial test: p=0.009), while vehicle treatment did not significantly alter the within-group proportion (n=2 out of 7, binomial test: p=0.331). Interestingly, treatment condition did not significantly affect other performance measures between groups, including the shape of the acquisition curve and final performance level at asymptote (Fig. S1a), or the total number of bar press responses during the Memory Test two days later (Fig. S1b). Therefore, the primary behavioral effect of HDAC3 inhibition was to shift within-group variability towards frequency-specific memory. Furthermore, a direct group comparison of ranked distributions of animals by the specificity of memory just reaches a statistical distinction between RGFP966 and vehicle-treated animals (Mann-Whitney one-tailed U-test_(7,6)_=8.0; p=0.050), which supports HDAC3 as a mechanism that drives memory in one direction towards specificity along a natural continuum of individual variability in memory. The link between HDAC3 and memory specificity revealed behaviorally is likely due to its role to promote the consolidation of learning experiences via frequency-specific auditory system neuroplasticity^24,35^.

**Fig. 2.**
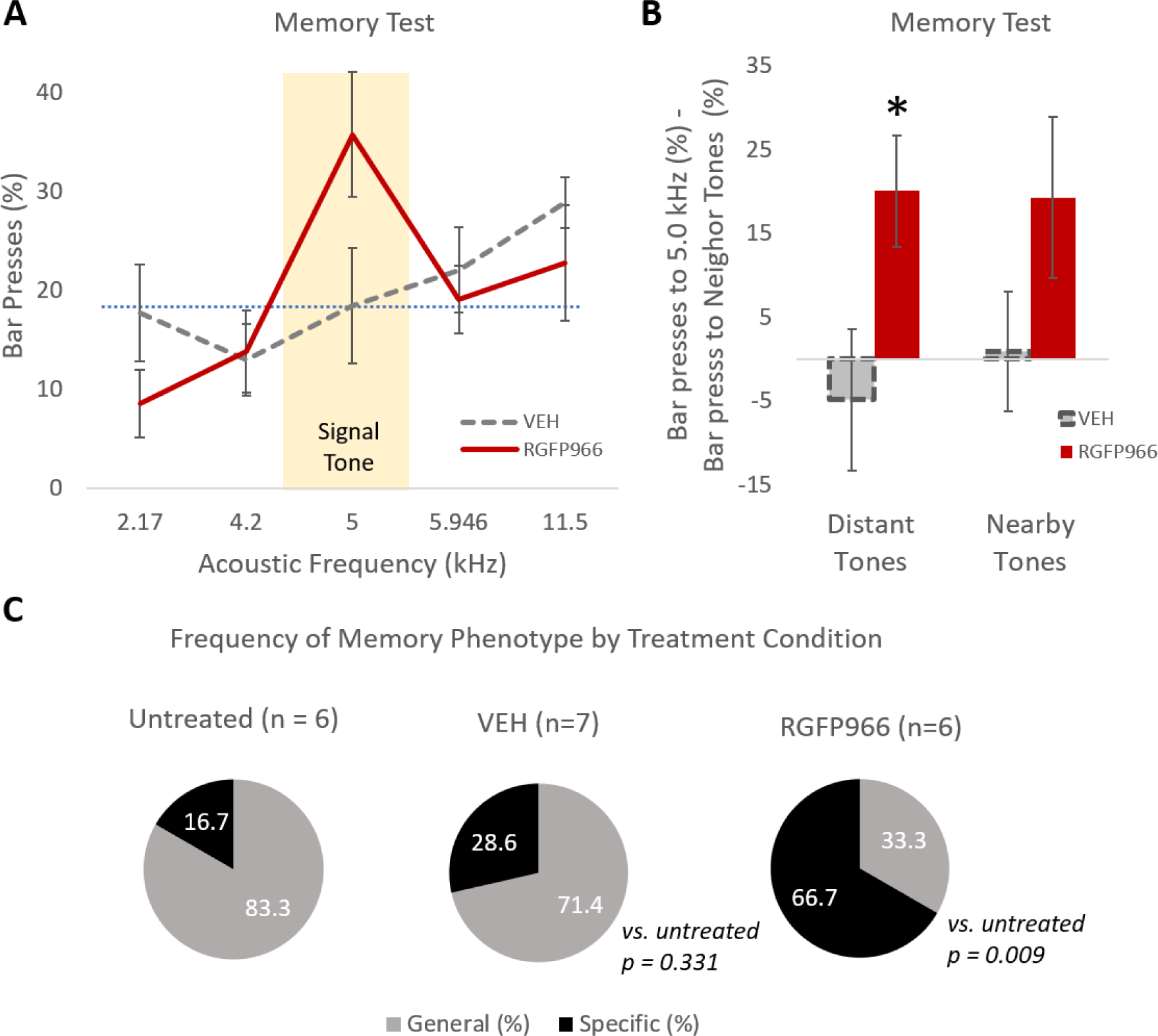
Post-training treatment with HDAC3 inhibitor RGF966 promotes frequency-specific memory. (A) RGFP966-treated rats exhibit a frequency-specific response distribution peaking at the signal tone frequency, while vehicle-treated rats exhibit a shallow response gradient. (B) Quantifying the shape of the response distribution using relative measures of responding to the signal tone vs. other test tone frequencies reveals that RGFP966-treated animals behaviorally discriminate. They respond to the signal tone more than both distant (far) tones (*left*) and nearby tones (*right*). Vehicle-treated rats do not discriminate, responding equally to signal tone vs. nearby or distant tones. (C) RGFP966 treatment significantly increases the proportion of individuals with frequency-specific memory type, compared with untreated individuals. Vehicle treatment does not alter the distribution of memory phenotype. All error bars represent +/- SEM. *p<0.05

Analyses of untreated animals in Fig. 1 identified a putative function of auditory system plasticity for memory specificity. Here, we sought to determine whether under conditions of pharmacological treatment, group differences in memory specificity revealed behaviorally between RGFP66-vs. vehicle-treated animals would mimic natural learning-induced individual variability in auditory system plasticity.

HDAC3 inhibition has known sequelae in learning-induced plasticity at the level of the primary auditory cortex (A1)^24,35^. Thus, electrophysiological recordings in A1 were obtained outside of the training context, and following the Memory Test were to identify forms of A1 plasticity from the treated groups (RGFP966 vs. vehicle) to compare to a separate group of 5 naïve rats. Electrophysiological data were analyzed according to the characteristic frequency of each site to parallel behavioral analyses by pooling neural data *near* the signal tone frequency (Fig. 3c) separately from sites tuned *far* from the signal tone frequency (Fig. 3d). Cortical receptive fields were analyzed for tuning bandwidth (BW, with respect to the breadth of frequency responsiveness in octaves) and response threshold (with respect to sound-evoked level in decibels). In sites tuned near the signal tone frequency (within one-third of an octave), one-way ANOVA revealed a significant difference in tuning bandwidth between groups (see note 1). Post hoc Holm-Bonferroni-corrected t-tests showed that RGFP966-treated animals had significantly narrower bandwidth than both naïve and vehicle-treated animals at each sound level tested above threshold (i.e., 10 dBs above threshold, BW10; and so on for BW20, BW30, BW40), while there were no significant differences between naïve or vehicle-treated animals at any bandwidth. Cortical sound-evoked response threshold did not differ among groups (naïve: n=28, M=21.63, SE=2.036; veh: n=44, M=22.39, SE=1.794; RGFP966: n=23, M=16.78, SE=2.543; one-way ANOVA: F(2,92)=1.6732, p=0.193).

**Fig. 3.**
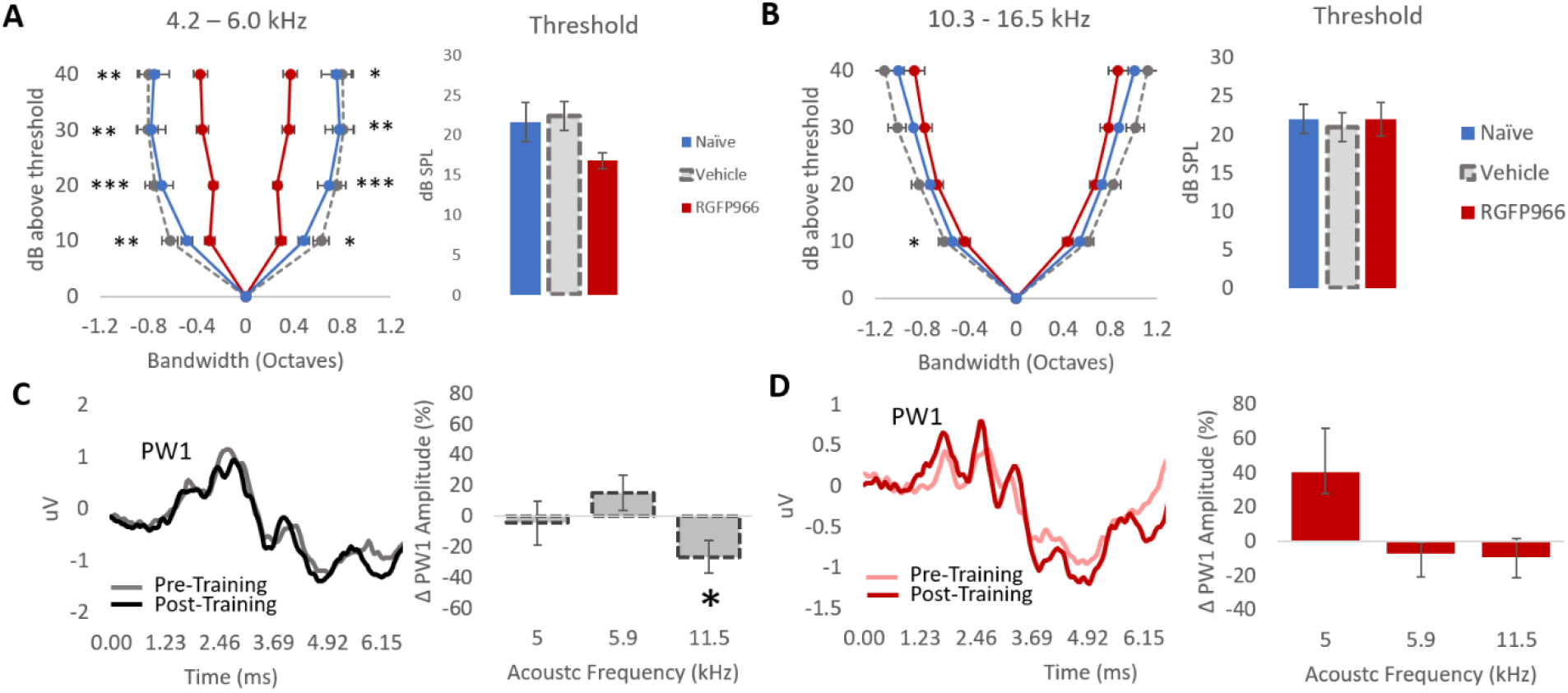
HDAC3 inhibition promotes learning-induced auditory system plasticity that is signal-specific. Panels represent sound-evoked neural responses from the auditory cortical (A,B) and auditory brainstem response (C,D) recordings. (A) Among auditory cortical sites tuned near the signal tone frequency (+/- 1/3 octave), RGFP966-treated animals showed significantly narrower tuning bandwidth at every sound level than vehicle-treated and naïve animals. Bandwidth among vehicle-treated and naïve animals did not differ. There were no group differences in response threshold (dB SPL). (B) Tuning bandwidth between-groups was more similar among auditory cortical sites tuned far away from the signal tone frequency (+1.03-1.33 octaves); RGF966-treated rats only had narrower BW10 than vehicle-treated rats (but were the same as naïve). (C) *Left*, Representative signal-tone evoked ABR traces from a single (out of 7) vehicle-treated subject recorded before (“pre-training,” gray) and after (“post-training,” black). *Right*, Quantification of learning-induced PW1 amplitude changes in ABRs evoked by the 5.0 kHz signal tone, as well a near (5.946 kHz) and far (11.5 kHz) neighbor frequency. There are no significant amplitude changes in 5.0 or 5.946 kHz evoked PW1, but a significant amplitude decrease in 11.5 kHz evoked PW1. (D) *Left*, Representative signal-tone evoked ABR traces from a single RGFP966-treated subject (out of 4) recorded before (gray) and after (black) training. *Right*, Quantification of learning-induced PW1 amplitude changes in ABRs evoked by the 5.0, 5.946, and 11.5 kHz, though none reached statistical significance. All error bars represent +/- SEM. *p<0.05 **p<0.01 ***p<0.001. In (A, B), asterisks on the left represent comparisons between vehicle and RGFP966; on the right, naïve vs. RGFP966. No significant differences were found between naïve and vehicle groups.

In recording sites tuned over an octave away from the signal tone frequency, there were no significant overall group differences in tuning bandwidth (see note 2), except BW10 (posthoc Holms-Bonferroni corrected t-tests show RGFP966-treated animals had narrower bandwidth than vehicle-treated animals, but not naïve animals, with no other group differences: Naïve vs. veh: t(107)=-1.097, p=0.275; naïve vs. RGFP966: t(99)=-1.671, p=0.218; veh vs. RGFP966: t(89)=2.522, p=0.039). Again, response threshold did not differ among groups (Naïve: n=53, M=22.01, SE=1.877; veh: n=62, M=21.65, SE=1.817; RGFP966: n=41, M=19.59, SE=2.309; one-way ANOVA: F(2,153)=0.040, p=0.960). These findings again support HDAC3 as a mechanism that drives neural substrates of memory in a direction towards specificity along a natural continuum of individual variability.

To determine whether HDAC3 effects also mimic subcortical neural substrates of frequency-specific memory, ABRs were recorded before and after training in the same groups of treated animals to quantify learning-induced change in PW1 amplitude to: the signal tone (5.0 kHz), a *nearby* tone frequency (5.946 kHz), or a *distant* tone frequency, over an octave away from the signal (11.5 kHz). Among vehicle-treated animals, there was no significant change in PW1 amplitude evoked the 5.0 kHz signal tone (M=-4.4738, SE=14.336, one sample t-test: t(6)=- 0.3314, p=0.766), or by the 5.946 kHz *near* tone (M=14.848, SE=11.568, one sample t-test: t(6)=1.283, p=0.246), while there was a small but significant amplitude change in the opposite direction (decrease) in PW1 amplitude evoked by the *far* 11.5 kHz tone (M=-26.738, SE=10.679, one sample t-test: t(6)=-2.503, p=0.046) (Fig. 3c). Among RGFP966-treated animals however, there was a notable increase in PW1 amplitude evoked by the 5.0 kHz signal tone, though not statistically significant (M=40.239, SE=25.809, one sample t-test: t(3)=1.559, p=0.218). RGFP966-treated animals also did not exhibit ABR amplitude changes evoked by either the 5.946 kHz tone (M=-7.056, SE=6.351, one sample t-test: t(3)=-1.110, p=0.346) or the 11.5 kHz tone in either direction (M=-9.4073, SE=10.825, one sample t-test: t(3)=-0.869, p=0.448). However, the negative findings of group treatments were based on ignoring the actual frequency-specificity of individual subject memory, which may have masked significant relationships that exist at the level of the individual subjects. Insofar as individualized analyses could be performed without compromising group-based evaluations, the same data were re-analyzed. Re-grouping the ABR data by “specific” (n=5/13) vs. “general” (n=6/13) memory phenotype (rather than by treatment) revealed greater signal-specific PW1 amplitude increases as individuals exhibit more frequency-specific memory behaviorally (Fig. S3c). Interestingly, re-grouping the cortical data also by “specific” (n=6/13) vs. “general” (n=7/13) memory phenotype (i.e., regardless of treatment condition) revealed the same significant brain-behavior relationship reported above: individuals with frequency-specific memory exhibited a cortical narrowing in tuning bandwidth in sites tuned only near the signal frequency, with no significant differences in threshold (Fig. S3a,b). Therefore, though HDAC3 inhibition via RGFP966 does promote frequency-specific memory^24,35^ (Fig. 2), this functional outcome may depend not just on the intervention alone, but on the ability of that intervention to facilitate the appropriate form, locus, and magnitude of auditory system plasticity that together provide sufficient neural substrates for the specificity of memory to reveal behaviorally. In the present case, memory specificity appears to require both cortical and subcortical substrates.

Thus far, the findings reveal that (1) treatment with HDAC3-inhibitor RGFP966 drives individual differences towards a frequency-specific memory phenotype, and (2) that frequency-specific memory is associated with forms of signal-specific plasticity at multiple levels of the auditory system. As such, intervention with RGFP966 treatment resulted in an opportunity to determine brain-behavior relationships to better understand the function of auditory system plasticity for auditory memory specificity. To capitalize on this opportunity, individual subjects were used to determine whether magnitude of effects in the forms of plasticity identified (both subcortically, as in Fig. 1; and cortically, as in Fig. 3) would reflect magnitude effects in the frequency-specificity of learned behavior.

Replicating the relationship observed in untreated animals (Fig. 1d), there was a significant positive correlation between the change in signal-tone evoked PW1 amplitude and the proportion of responses to the signal tone during Memory Test in treated animals (*r=0.890, p=0.0002)*. The greater the amplitude gain, the greater the proportion of bar-presses to the signal frequency (Fig. 4b). Notably, there was no relationship between pre-training PW1 amplitude and subsequent responses at Memory Test (*r=0.137, p=0.665)*, which supports that all reported relationships are learning-induced (Fig. 4a). To quantify memory specificity at the individual level, two response contrast measures were derived between pairs of neighboring tone frequencies for each subject: (1) for behavioral contrast, as the difference in bar-presses to the signal tone vs. near or distant neighboring tone, and (2) for neural contrast, as the difference in PW1 amplitude changes (i.e., before-*minus* after-training) evoked by the signal tone vs. neighboring tone. Figure 4c shows a significant positive correlation discovered between subcortical neural and behavioral contrasts for the 5.0 kHz signal tone and the near neighbor tone 5.946 kHz (*r=0.727, p=0.011)* as well as its distant neighbor tone 11.5 kHz (*r=0.696, p=0.017*) (Fig. 4d): greater neural contrast predicts greater behavioral contrast. Further, there was a significant cortical correlate of frequency-specific behavioral responding: Cortical tuning bandwidth negatively correlated with the proportion of bar presses to the signal tone during the Memory Test *(r=-0.668, p=0.017)* (Fig. 4e). Thus, narrower signal-specific cortical tuning also predicted greater behavioral signal contrast.

**Figure 4.**
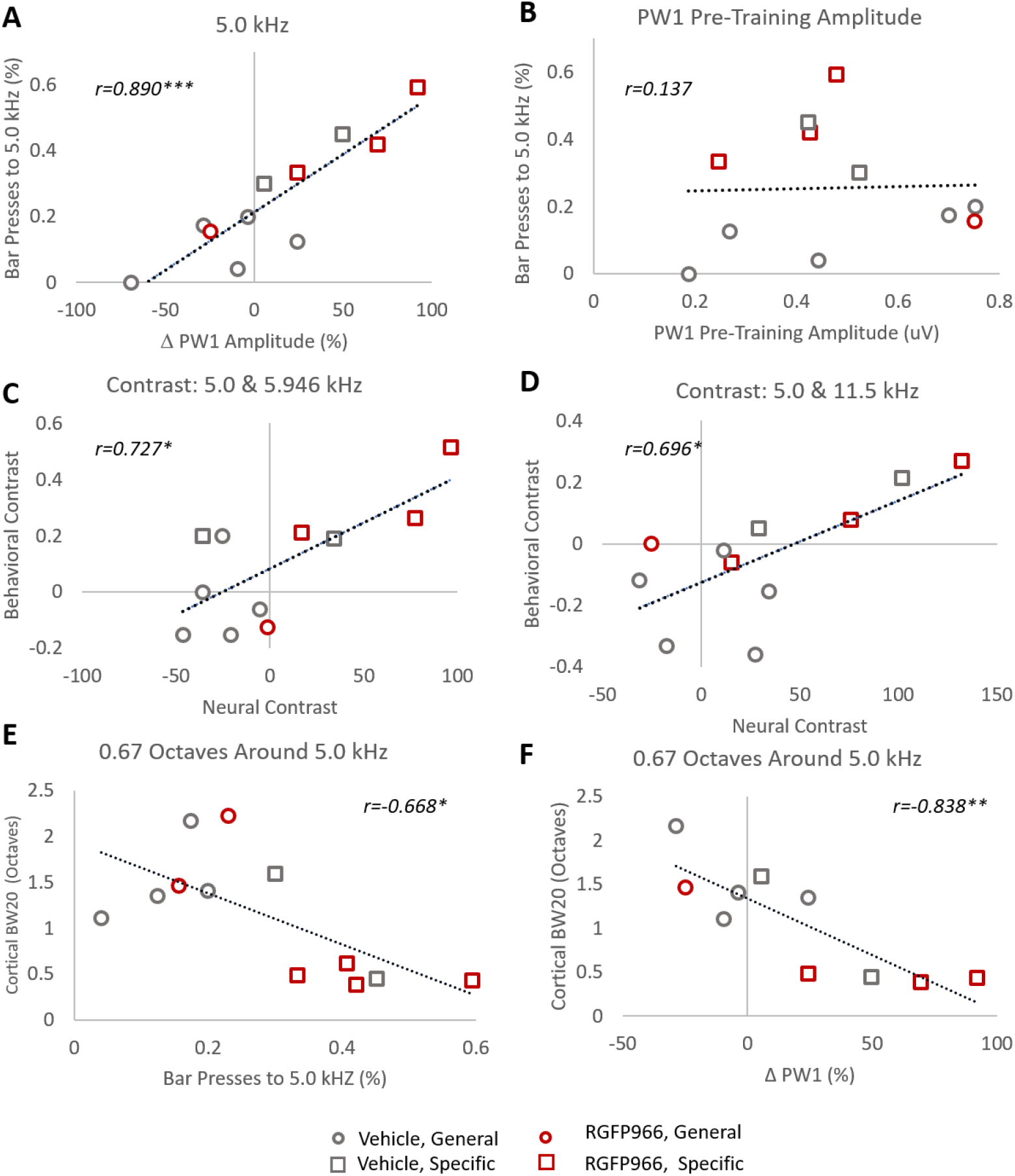
Coordinated forms of auditory system plasticity are correlated with the frequency-specificity of auditory memory. (A) Greater amplitude gains in signal-tone evoked PW1 predict a greater percent of total responses to the signal tone frequency at Memory Test. (B) Pre-training PW1 amplitude has no relationship with subsequent memory specificity. (C) Greater neural contrast between the signal tone and a near neighbor tone (as measured by PW1 amplitude changes) predicts greater behavioral contrast (measured by difference in percent of bar presses) to that same pair of tones. (D) Greater neural contrast between the signal tone and a distant neighbor tone also predicts greater behavioral contrast among that pair of tones. (E) Narrower auditory cortical tuning bandwidth (BW20) for sites tuned near the signal tone correlates with a greater percentage of responses to the signal tone. (F) Narrower auditory cortical tuning bandwidth (BW20) for sites tuned near the signal tone is predicted by greater amplitude gains in signal tone-evoked PW1. *p<0.05 **p<0.01 ***p<0.001

We linked the two analyses together to here report for the first time a putative connection between subcortical and cortical forms of plasticity accompanying the formation of signal-specific memory. There is a significant correlation between the amplitude change of 5.0 kHz-evoked PW1 and signal-specific auditory cortical tuning bandwidth *(r=-0.838, p=0.0024)*: as cortical bandwidth decreases, PW1 amplitude increases (Fig. 4f). Thus, individual differences in two identified forms of learning-induced auditory neuroplasticity validated them both as coordinated substrates of memory’s acoustic specificity. In sum, these findings confirm and extend a hypothesized relationship between cortical plasticity, subcortical plasticity, and learned behavior.

## Discussion

We report three main findings. (1) Frequency-specific memory, whether *natural* or mediated by a HDAC3 pharmacological intervention, is associated with multiple forms of signal-specific auditory system plasticity that includes cortical and subcortical candidate substrates. (2) Inhibition of HDAC3 during early auditory learning of a single-tone task promotes a lasting frequency specific memory phenotype. (3) A three-way correlation between auditory cortical plasticity, auditory subcortical plasticity, and sound-specific behavior revealed by within-group individual differences validates neural candidates as substrates of memory revealed in behavior.

A three-way correlation between sound-specific cortical and subcortical plasticity, and behavior supports that learning can induce coordinated auditory system reorganization at multiple levels. When those forms of plasticity emerge in a signal-specific way for acoustic frequency, the learned behavior emerges with signal-specificity for acoustic frequency. Furthermore, the magnitude of learning-induced cortical changes explains the magnitude of amplitude changes even in the earliest peak of the ABR, thought to be generated as early as the auditory nerve^40,41^. Thus, coordination appears to span the highest to lowest levels of the auditory system, which is consistent with previous corticofugal interpretations of long-lasting experience-dependent plasticity in the human ABR^42^ and the brainstem’s frequency-following response (FFR)^43^. Indeed, the interaction between cortical and subcortical sound-evoked responses is not new^44^, which includes evidence of cortical re-organization of descending inputs after early auditory system damage^45^. The present findings relate these interactions to adult learning-dependent effects. Importantly, ABR plasticity that was detected outside of the training context after achieving high-levels of performance predicted *subsequent* specificity of behavioral performance in a completely novel situation, days later, at Memory Test. Therefore, noninvasive ABR neurophysiology has high translational potential to track and anticipate the effects of learning for sound-specific behavioral functions on an individual subject basis, which is relevant for the successful transfer of training experiences from a clinical setting to real-life contexts.

In addition to demonstrating a function of signal-specific auditory system plasticity for memory specificity, the findings confirm HDAC3 as a mechanism of long-term memory formation by promoting signal-specific auditory system plasticity, which now includes subcortical effects. Future research is necessary to determine whether the effects of systemic delivery of an HDAC3 inhibitor facilitates learning-induced neuroplasticity directly in both sensory subcortical and cortical areas, or whether its effects on the ABR are sequelae of primary effects in the cortex. In either case, the overall effect of HDAC3 inhibition appears to be a significant shift along a natural continuum of individual variability in memory formation towards specificity, maintaining the same characteristic neurophysiological substrates of auditory memory as untreated, “naturally”-learning subjects. Therefore, HDAC-targeted manipulations may be useful tools to study the links between neurophysiological and molecular substrates of long-term memory in the brain. Used in the auditory and other sensory systems, these tools could provide answers to how multidimensional memories are stored with multi-feature specificity and its consequence for future behavioral action.

A remaining open question is: *Where do individual differences come from?* The answer is beyond the scope of the current studies, but not beyond the scope of behavioral neuroscience, where much work is being done to identify key factors. Here, all animals were the same species, strain, sex, age, and all were experimentally naïve prior to training. Therefore, other putative factors related to the integrity of anatomical circuitry, efficiency of experience to drive neuromodulatory events, or the availability of molecular signaling factors could be examined. Recent evidence in adult and aging subjects points to medial temporal lobe function and anatomical integrity as a likely locus of effect for the specificity (vs. generalization) of episodic memory^46^, which may prove to interact with sensory cortical and subcortical substrates^47-50^. Regardless of the source of individual differences in the brain, or their environmental causes, the lesson learned from this work is that to harness plasticity mechanisms for adaptive goals, we must identify neurophysiological substrates in form and magnitude that match their desired functional outcomes. Since the individual is of prime importance in a clinical setting, these issues remain a critical subject for future behavioral neuroscientific research.

## Materials and Methods

### Subjects

A total of 19 adult male Sprague-Dawley rats (275-300 g on arrival; Charles River Laboratories, Wilmington MA) were used (n= 6 untreated; n = 13 treated) in behavioral and electrophysiological procedures. Additionally, 5 naïve adult males were used for cortical electrophysiological recordings. All animals were individually housed in a colony room with a 12-hour light/dark cycle. Throughout behavioral procedures, rats were water-restricted, with daily supplements provided to maintain at ∼85% free-drinking weight^24,35^. All procedures were approved and conducted in accordance with guidelines by the Institutional Animal Care and Use Committee at Rutgers, The State University of New Jersey.

### Behavioral Procedures and Analysis

All behavioral sessions were conducted in instrumental conditioning chambers within a sound-attenuated box. All subjects initially learned how to press a lever for water reward in five ∼45-minute bar-press shaping sessions. Next, all rats underwent *tone-reward training*, a single tone detection task, in which they learned to associate a 5.0 kHz signal tone with the operant reward. Responses in the presence of the signal tone were rewarded, while responses during the intertrial interval (ITI) triggered a visual error signal and a time-out the extended that time until the next tone trial. All rats were trained to performance criteria, where on average 70% of bar presses occurred in the presence of the signal tone for 2 consecutive days (average training sessions for all subjects: n=19, M=12.10 +/-1.5). A two-way ANOVA was used to compare group performance on the first two tone-reward training sessions and the final two tone-reward training sessions.

Forty-eight hours following the final tone-reward training session, rats were tested in the Memory Test to determine their memory specificity for the signal tone frequency. In the Memory Test, rats were presented with the 5.0 kHz signal tone, as well as 4 novel tone frequencies representing “nearby” neighbors (+/- 0.25 octaves) to the signal tone (5.946 & 4.2 kHz) and “distant” neighbors (+/- 1.20 octaves) to the signal tone (11.5 & 2.17 kHz). No responses were reinforced. The distribution of bar presses among the test tone frequencies was used to determine the shape of the frequency generalization gradient (Shang et al., 2019). To quantify memory specificity for the signal tone, contrast measures of relative to response to the signal tone, vs. novel tones, were calculated as follows: (1) Percent of responses to signal tone – (average percent of response to distant tones) and (2) Percent of responses to signal tone – (average percent of responses to nearby tones). Positive values indicate greater responding to the signal tone than novel tones. Single-sample t-tests were used to determine whether contrast scores were significantly different than 0. To determine memory phenotype, we defined animals with frequency-specific memory as those who had *positive* contrast values for both distant and nearby tones (relative to the signal tone). A binomial test was used to determine the categorical frequency of memory phenotype by treatment condition, compared to untreated subjects. A one-tailed Mann-Whitney U test was used to determine differences in rank-order distribution of memory specificity by treatment condition. For Pearson correlative data, behavioral contrast measures were also derived between the signal tone and a single neighbor to match the sound frequencies used in auditory brainstem response recordings.

### Pharmacological Inhibition of HDAC3

A pharmacological HDAC3 inhibitor RGFP966 was used alter molecular mechanisms of auditory memory formation induced by learning^35^. Rats in this treatment experiment were randomly assigned to either the RGFP966 (n=6) or vehicle (n=7) condition prior to tone reward-training. Rats received 3 consecutive days of post-session injections of RGFP966 (10 mg/kg, s.c.) or vehicle (equated for volume) on training days 2-4 (dose established^51^; and confirmed in auditory system function^35^). Post-training pharmacological treatment confines manipulation to the memory consolidation period, while avoiding potential performance effects based on perception, motivation or within-session learning. For the remainder of training sessions after day 4, all rats received post-session injections of saline (equated for volume) to ensure that any effect of the injection itself remained consistent throughout training until reaching performance asymptote.

### Auditory Brainstem Response Recordings and Analysis

Auditory brainstem responses (ABRs) were recorded twice in anesthetized rats (sodium pentobarbital, 50 mg/kg, i.p.) to determine learning-induced changes in subcortical sound processing: (1) 24 hours prior to the first tone-reward training session and (2) 24 hours following the final-tone reward training session. All recordings were made in a recording chamber completely separate from the training chamber and while the animal was anesthetized, which is a completely different state and context than that used in training. Moreover, animals in Stimulus presentation and neural response recordings were carried out using BioSig RZ software (TDT). Evoked potentials were recorded using a three-electrode configuration, with subdermal needle electrodes (1 kΩ) positioned at the midline along the head (recording), immediately below the left pinna (reference), and the midline on the back of the neck (ground) Sound stimuli were 60 dB SPL, 5ms pure-tones (2 ms cosine-gated rise/fall time) presented at 21 Hz to the left ear from a speaker positioned 4 cm away. Three tone frequencies (11.5, 5.946, and 5.0 kHz) were presented in a blocked format (512 stimuli per block). The averaged evoked response was used for analysis of the first positive peak (PW1) of the waveform. Custom Matlab© scripts were used to identify peaks within the waveform and derive the trough-to-peak amplitude (uV). Learning-induced amplitude changes were calculated as: (Post-training amplitude – pre-training amplitude)/pre-training amplitude)) * 100. Two-tailed single-sample t-tests were used to determine significant amplitude changes as a function of learning. For Pearson correlative data, neural contrast scores were calculated as: Learning-induced amplitude change in ABR evoked by the signal tone – Learning-induced amplitude change in ABR evoked by a neighbor tone. We were unable to obtain valid post-training for 2 subjects, resulting in the following group numbers: vehicle: n=7/7; RGFP966: n=4/6; general memory: n=6/7; specific memory: n=5/6.

### Auditory Cortical Recording Procedures and Analysis

To determine changes in the frequency-specificity of auditory cortical bandwidth tuning, electrophysiological recordings were obtained from anesthetized subjects (n=13) (sodium pentobarbital, 50mg/kg, i.p.) in an acute, terminal recording session 24-48 hours following the Memory Test. All recordings were made the same recording chamber as the was used to obtain ABRs, which was completely separate from the training chamber and in a completely different state and context than that used in training. Recordings were also obtained from a group of naïve rats (n=5). Recordings were performed inside a double-walled, sound attenuated room using a linear array (1 × 4) of parylene-coated microelectrodes (1-2 MΩ, 250 µm apart) targeted to the middle cortical layers (III-IV, 400-600 µm orthogonal to the cortical surface) of the right auditory cortex. Multiple penetrations were performed across the cortical surface (M=63.55 sites/animal, SE=3.62). Acoustic stimuli were presented to the left ear from a speaker positioned ∼10 cm from the ear. Sounds were 50 ms pure tones (1-9 ms cosine-gated rise/fall time) presented in a pseudorandom order (0.5-54.0 kHz in quarter-octave steps; 0-70 dB SPL in 10 dB steps; 5 repetitions) with a variable inter-stimulus interval an average of 700 +/- 100ms apart. Neural activity was amplified x1000 and digitized for subsequent off-line spike detection and analysis using custom Matlab© scripts. Recordings were bandpass filtered (0.3-3.0 kHz). Multiunit discharges were characterized using previously reported temporal and amplitude criteria^52^. Acceptable spikes were designated as waveforms with peaks separated by no more than 0.6 ms and with a threshold amplitude greater than 1.5 (for the positive peak) and less than 2.0 (for the negative peak) × RMS of 500 random traces from the same recording on the same microelectrode for each site. For each recording site, tone-evoked spike rate (spikes/s) were calculated by subtracting spontaneous spiking (40 ms window prior to tone onset) from evoked-spiking within a 40 ms response-onset window (6-46 ms after each tone onset). Responses greater than +/−1.0 SEM of the spontaneous spike rate were considered true evoked responses. Tone-evoked activity was used to construct frequency-response areas (FRAs) for each recording site, which reveal the mean sound-evoked activity to each frequency/sound level combination. The borders of each FRA were determined based on a threshold firing rate value determined by its spontaneous activity. Only evoked responses greater than the mean of pre-onset spontaneous activity were considered true sound-evoked responses. The outside border of each FRA was used to determine (1) response threshold, or the lowest sound level (dB SPL) that evokes a response, (2) characteristic frequency (CF), or the frequency to which the site responds most strongly (in spikes/s) at threshold sound level, and (3) tuning bandwidth, or the breadth of frequency responsivity (in octaves) as a function of dB above response threshold. Thus, bandwidth10 (BW10), BW20, BW30, and BW40 denote bandwidth 10, 20, 30, and 40 dB SPL above threshold sound level, respectively. In order to determine tuning plasticity as a function of acoustic frequency, bandwidth data was sorted by CF to create two frequency bins: (1) sites tuned near (within +/− 1/3 octave) the 5.0 kHz signal tone frequency (4.2-6.0 kHz) and (2) sites tuned far (between 1.04 and 1.70 octaves) away from the 5.0 kHz signal tone frequency (10.3-16.5 kHz). For group analysis of bandwidth, individual recording sites were treated as individual observations (naïve: n=5 subjects/26 recordings sites near 5.0 kHz/50 recording sites far from 5.0 kHz; vehicle: n=7 subjects/44 sites near 5.0 kHz/59 sites far from 5.0 kHz; RGFP966: n=6 subjects/23 sites near 5.0 kHz/41 sites far from 5.0 kHz). Differences in tuning bandwidth within a frequency bin was compared among conditions (vehicle, RGFP966, and naïve) using one-way ANOVA. Pairwise comparisons were made with Holm-Bonferroni corrected two-tailed t-tests. Corrected p-values are reported. For Pearson correlative data, an average bandwidth score for BW20 was computed for each individual. One outlier belonging to the vehicle/frequency-general memory groups was excluded from analysis justified by >3 times the mean Cook’s distance was excluded from the cortical correlations.

*The data that support the findings of this study are available from the corresponding author upon reasonable request.*

## Acknowledgments

The authors word like to thank Ms. Andrea Shang for collecting auditory cortical data from naïve subjects, Ms. Alyssa Rodriguez for assisting in behavioral training, and the CLEF Lab personnel for their support.

## Author contributions

E.K.R. and K.M.B. designed the experiment. E.K.R. collected the data. E.K.R. and K.M.B. analyzed the data and wrote the manuscript. E.K.R. prepared the figures.

## Data and materials availability

All data, code, and materials used in the analysis will be made available to any researcher for purposes of reproducing or extending the analysis at request.

## Footnotes

1 Complete statistical results: **BW10**-naïve: n=26, M=0.96, SE=0.092; veh: n=44, M=1.24, SE=0.126; RGFP966: n=23, M=0.586, SE=0.084; one-way ANOVA: F(2,90)=7.602, p=0.0008; Holms-Bonferroni corrected two-tailed t-test: naïve vs. veh: t(68)=-1.612, p=0.114; naïve vs. RGFP966:t(47)=2.9259, p=0.014; veh vs. RGFP966:t(65)=3.558, p=0.0021; **BW20-** naïve: M=1.38, SE=0.188; veh: M=1.50, SE=0.162; RGFP966: M=0.52, SE=0.073; one-way ANOVA: F(2,90)=9.193, p=0.0002; Holms-Bonferroni corrected two-tailed t-test: naïve vs. veh: t(68)=-0.445, p=0.657; naïve vs. RGFP966: t(47)=4.048, p=0.0002; veh vs. RGFP966: t(65)=4.255, p=0.00021; **BW30 –** naïve: M=1.55, SE=0.236; veh: M=1.61, SE=0.166; RGFP966: M=0.71, SE=0.098; one-way ANOVA: F(2,90)=6.366, p=0.0025; Holms-Bonferroni corrected two-tailed t-test: naïve vs. veh: t(68) =- 0.2062, p=0.837; naïve vs. RGFP966: t(47)=3.122, p=0.006l veh vs. RGFP966: t(65)=3.74, p=0.0013; **BW40 –** naïve: M=1.50, SE=0.242; veh: M=1.60, SE=0.173; RGFP966: M=0.74, SE=0.111; one-way ANOVA: F(2,90)=5.251, p=0.0069. Holms-Bonferroni corrected two-tailed t-test:naïve vs. veh: t(68)=-0.328, p=0.743; naïve vs. RGFP966: t(47)=2.734, p=0.0174; veh vs. RGFP966: t(65)=3.376, p=0.0036

2 Complete statistical results: **BW10-** naïve: n=50, M=1.08, SE=0.082; veh: n=59, M=1.22, SE=0.088; RGFP966: n=41, M=0.87, SE=0.101; one-way ANOVA: F(2,147)=3.468, p=0.033; **BW20-** naïve: M=1.47, SE=0.211; veh: M=1.62, SE=0.119; RGFP966: M=1.34, SE=0.109; one-way ANOVA: F(2,147)=1.585, p=0.211; **BW30**-naïve: M=1.74, SE=0.119; veh: M=1.96, SE=0.148; RGFP966: M=1.58, SE=0.145; one-way ANOVA: F(2,147)=1.829, p=0.164; **BW40 –** naïve: M=2.01, SE=0.155; veh: M=2.18, SE=0.175; RGFP966: M=1.72, SE=0.175; one-way ANOVA: F(2,147)=1.806, p=0.167

