## Supplementary Materials for "Precise Memory is Predicted by Learning-Induced Sensory System Neurophysiological Plasticity"

**Text:**

Figure S1 presents performance metrics for RGFP966- and vehicle-treated animals (Vehicle: n=7, first two training sessions: M=10.07, SE=1.28. last two training sessions: M=78, SE= 3.26; RGFP966: n=6, first two training sessions: M=9.85, SE=1.38, last two training sessions: M=79.5, SE=2.88) Two-way ANOVA revealed a main effect for training session, such that performance accuracy increased with training ($F(1,22)=113.504$, $p<0.001$). However, there was no main effect for treatment condition ($F(1,22)=1.069$, $p=0.312$) and no significant training session x treatment interaction ($F(1,22)=0.933$, $p=0.33$), demonstrating that treatment with the HDAC3-inhibitor, RGFP966, did not significantly alter performance during tone reward training (Fig. S1a,b). Further, RGFP966 treatment did not alter the total number of bar presses made during the Memory Test (Fig. S1b), either in the presence of tones (Vehicle: M = 29.714, SE = 4.079; RGFP966: M = 34.333, SE = 1.855; two-tailed independent t-test: $t(6,5) = -0.973$, $p = 0.351$ ) or during the overall session (Vehicle: M = 38.286, SE = 4.47; RGFP966: M = 40.167, SE = 2.74; two-tailed independent t-test: $t(2,11)= -0.343$, $p = 0.737$). Therefore, a comparison of behavioral responses in the Memory Test at the conclusion of training cannot be attributed to the success or level of performance achieved. Rather, differences are attributed to the specificity of memory for frequency.

Figures S2 presents behavioral performance during the Memory Test when individual animals are grouped by their memory phenotype as “specific” or “general” (as described in the main text), rather than by pharmacological treatment condition. Rats with frequency-specific memory made a significantly greater percent of responses to the signal tone compared to distant tones (M=25.65, SE=5.27; two-tailed single sample t-test: $t(5)=4.862$, $p=0.004$) and compared to nearby tones (M=28.89, SE=5.88; two-tailed single sample t-test: $t(5)=4.917$, $p=0.004$) (Fig. S2b). This indicates their behaviorally ability to remember and discriminate the signal from non-signal tones. In contrast, rats with frequency-general memory did not behaviorally discriminate the signal tone from either the distant tones (M=-9.62, SE=5.93; two-tailed single sample t-test: $t(6)=-1.624$, $p=0.155$) or the nearby tones (M=-7.3, SE=4.23; two-tailed single sample t-test: $t(6)=-1.725$, $p=0.135$) (Fig. S2b). Despite differences in memory specificity, there were no group differences in the number of bar presses made during the Memory Test, both in the presence of tones (General: M=31, SE=3.559; Specific: M=32.83, SE=3.30; two-tailed independent sample t-test: $t(2,11)=-0.372$, $p=0.716$) and overall (General: M=40.57, SE=3.63; Specific: M=37.5, SE=4.03; two-tailed independent samples t-test: $t(2,11)=-0.566$, $p=0.582$). Since the denominator remains the same, the use of the distribution of all bar-press responses to the various Memory Test tone frequencies as a proportional value is valid to examine and compare the specificity of memory specificity between animals and groups.

Figure S3 presents auditory cortical (A,B) and auditory brainstem response (C) recording data when RGFP966- and vehicle-treated animals are grouped by memory phenotype (as in Fig.

S2). Each recording site was characterized by its characteristic frequency (CF; the frequency that evoked a response at the lowest sound level). Among cortical sites with CFs that were tuned near the signal tone frequency (within 1/3rd of an octave; Fig. S3a), animals with frequency-specific memories had significantly narrower bandwidth at all sound levels (expressed as dB-steps above threshold, i.e., the sound level at which a tone could evoke a neural response), compared to animals with general memories. There was also a significance difference in the breadth of tuning between animals with frequency-specific memories at BW20, BW30, and BW40 compared to naïve animals (**BW10** – naïve: n = 26, M = 0.96, SE = 0.092; general: n = 38, M = 1.28, SE = 0.138; specific: n = 29, M = 0.68, SE = 0.097; one-way ANOVA: F(2, 90) = 6.540, p = 0.002; Holms-Bonferroni corrected two-tailed independent samples t-test: naïve vs. general: t(62) = -1.730, p = 0.088; naïve vs. specific t(53) = 2.019, p = 0.096; general vs. specific: t(65) = 3.289, p = 0.004; **BW20** – naïve: M = 1.38, SE = 0.188; general: M = 1.53, SE = 0.177; specific: M = 0.69, SE = 0.114; one-way ANOVA: F(2,90) = 7.338, p = 0.001; Holms-Bonferroni corrected two-tailed independent samples t-test: naïve vs. general: t(62) = -0.547, p = 0.586; naïve vs. specific t(53) = 3.243, p = 0.004; general vs. specific: t(65) = 3.745, p = 0.001; **BW30** – naïve: M = 1.55, SE = 0.236; general: M = 1.70, SE = 0.180; specific: M = 0.78, SE = 0.112; one-way ANOVA: F(2,90) = 7.204, p = 0.002; Holms-Bonferroni corrected two-tailed independent samples t-test: naïve vs. general: t(62) = -0.494, p = 0.622; naïve vs. specific t(53) = 3.024, p = 0.007; general vs. specific: t(65) = 3.977, p = 0.0005; **BW40** – naïve: M = 1.50, SE = 0.242; general: M = 1.63, SE = 0.242; specific: M = 0.88, SE = 0.135; one-way ANOVA: F(2,90) = 4.415, p = 0.014. Holms-Bonferroni corrected two-tailed independent samples t-test: naïve vs. general: t(62) = -0.424, p = 0.672; naïve vs. specific t(53) = 2.319, p = 0.048; general vs. specific: t(65) = 3.063, p = 0.009). There were no group differences in cortical response threshold (naïve: n = 28, M = 21.63, SE = 2.036; general: n = 38, M = 23.34, SE = 2.023; specific: n = 29, M = 16.71, SE = 2.034; one-way ANOVA: F(2,92) = 2.530, p = 0.085).

Among cortical sites with CFs tuned farther away, over an octave away from the signal tone (Fig. S3b), there were no group differences in bandwidth (**BW10** – naïve: n = 50, M = 1.08, SE = 0.082; general: n = 54, M = 1.19, SE = 0.091; specific: n = 46, M = 0.94, SE = 0.101; one-way ANOVA: F(2,147) = 1.867, p = 0.158; **BW20** – naïve: M = 1.47, SE = 0.211; general: M = 1.53, SE = 0.129; specific: M = 1.52, SE = 0.108; one-way ANOVA: F(2,147) = 0.944, p = 0.944; **BW30** – naïve: M = 1.74, SE = 0.119; general: M = 1.82, SE = 0.158; specific: M = 1.83, SE = 0.143; one-way ANOVA: F(2,147) = 0.090, p = 0.913; **BW40** – naïve: M = 2.01, SE = 0.155; general: M = 2.03, SE = 0.179; specific: M = 1.99, SE = 0.181; one-way ANOVA: F(2,147) = 0.003, p = 0.996) or response threshold (naïve: n = 53, M = 22.01, SE = 1.877; general: n = 57, M = 21.22, SE = 2.099; specific: n = 46, M = 20.34, SE = 1.871; one-way ANOVA: F(2,153) = 0.169, p = 0.844). Together, these data support that signal-specific memory for frequency (revealed behaviorally) is supported by signal-specific cortical plasticity that narrows tuning bandwidths, but only in sites tuned near (within 1/3rd octave) of the trained signal frequency.

At the subcortical level, an analysis of learning-induced changes in PW1 amplitude of the auditory brainstem response (ABR) likewise revealed learning-induced neural signal-specificity in animals with signal-specific memory. There was a signal-specific increase in the amplitude of response to the 5.0 kHz signal frequency in animals who later showed frequency-specific memory behaviorally (5.0 kHz – M=48.229, SE = 15.387; t(4) = 3.134, p = 0.035; 5.946 kHz- M = 10.361, SE = 8.809; t(4) = 1.176, p = 0.306; 11.5 kHz- M = -22.707, SE = 11.052; t(4) = -2.054, p = 0.109). No significant amplitude changes were detected in animals with frequency-general memories (5.0 kHz – M=-18.584, SE = 12.702; t(5) = -1.463, p = 0.203; 5.946 kHz- M = 3.981, SE = 13.738; t(5)

= 0.289, p = 0.783; 11.5 kHz- M = -18.543, SE = 12.206; t(5) = -1.519, p = 0.189). These data support that subcortical plasticity is in line with cortical plasticity substrates of behavioral signal-specific memory.

Table S1 displays Pearson r-values for correlations in brain vs. behavior or brain vs. brain measures in the overall group, as well as in subgroups that include pharmacological treatment conditions, and memory phenotypes. The subgroup correlations typically mimic those observed in the overall group. However, the general memory subgroup exhibits some of the weakest correlation strengths, vs. all other subgroups, in particular for correlations involving subcortical neural plasticity. This suggests that animals with general memories may be categorically different than animals with specific memories. Further, the general memory group is unique in that, unlike all other groups, it exhibits a *positive* correlation between cortical bandwidth and the percentage of responses to the 5.0 kHz signal tone at Memory Test, which is consistent with previous studies on sensory substrates of generalized memory in multiple modalities that indicate broadened tuning[53-57]. Together, this suggests that the forms of signal-specific auditory system plasticity reported here are particularly significant substrates of memory specificity, while other forms, brain regions or mechanisms may contribute to the generalization of memory.

**Figures:**

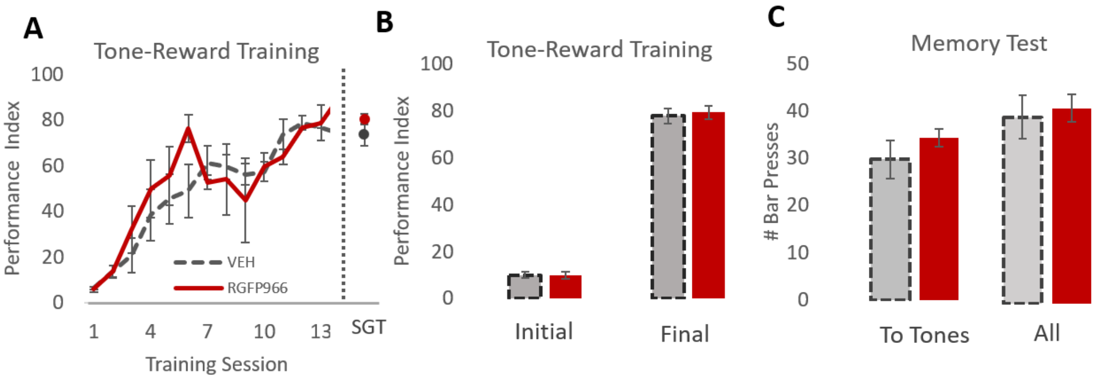


**Fig. S1.** *Treatment with RGFP966 does not alter acquisition performance or number of responses during memory test.* (A) RGFP966-treated rats learned to perform the task to an equal level of final performance as vehicle-treated rats. (B)Both RGFP966- and vehicle-treated group performance indices increased over the course of training, with no between-group differences in initial or final level of performance as assessed over the first two vs. final two training sessions. (C) During the memory test, all rats, whether RGF966- or vehicle-treated, made equal numbers of bar presses, both in the presence of tones and throughout the session overall. Errors bars represents +/- SEM. ***p<.0001

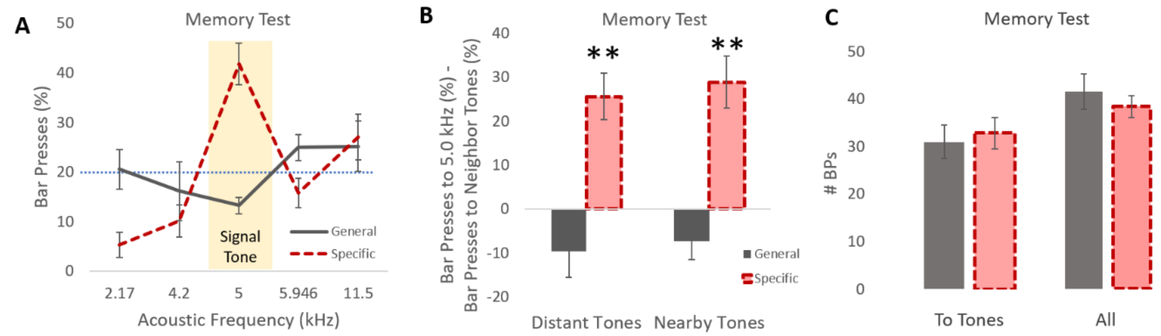


**Fig. S2.** *Analysis of behavioral performance with subjects grouped by memory phenotype*. (A) Rats with frequency-specific memory exhibited a frequency-specific response distribution, while those with frequency-specific memory exhibited a shallower response gradient. (B) Quantifying the shape of the response distribution using relative measures of responding to the signal tone vs. other test tone frequencies reveals that animals with frequency-specific memory behaviorally discriminate neighboring sound-frequencies. They respond to the signal tone more than both distant (far) tones (*left*) and nearby tones (*right*). Animals with frequency-general memory do not discriminate, responding equally to signal tone vs. nearby or distant tones. (C) Despite differences in memory specificity, rats with frequency-specific and -general memories made equal numbers of bar presses during the Memory Test, both to tones specifically (*left)* and throughout the test session overall *(right)*. Error bars represent +/- SEM. ***p<.001

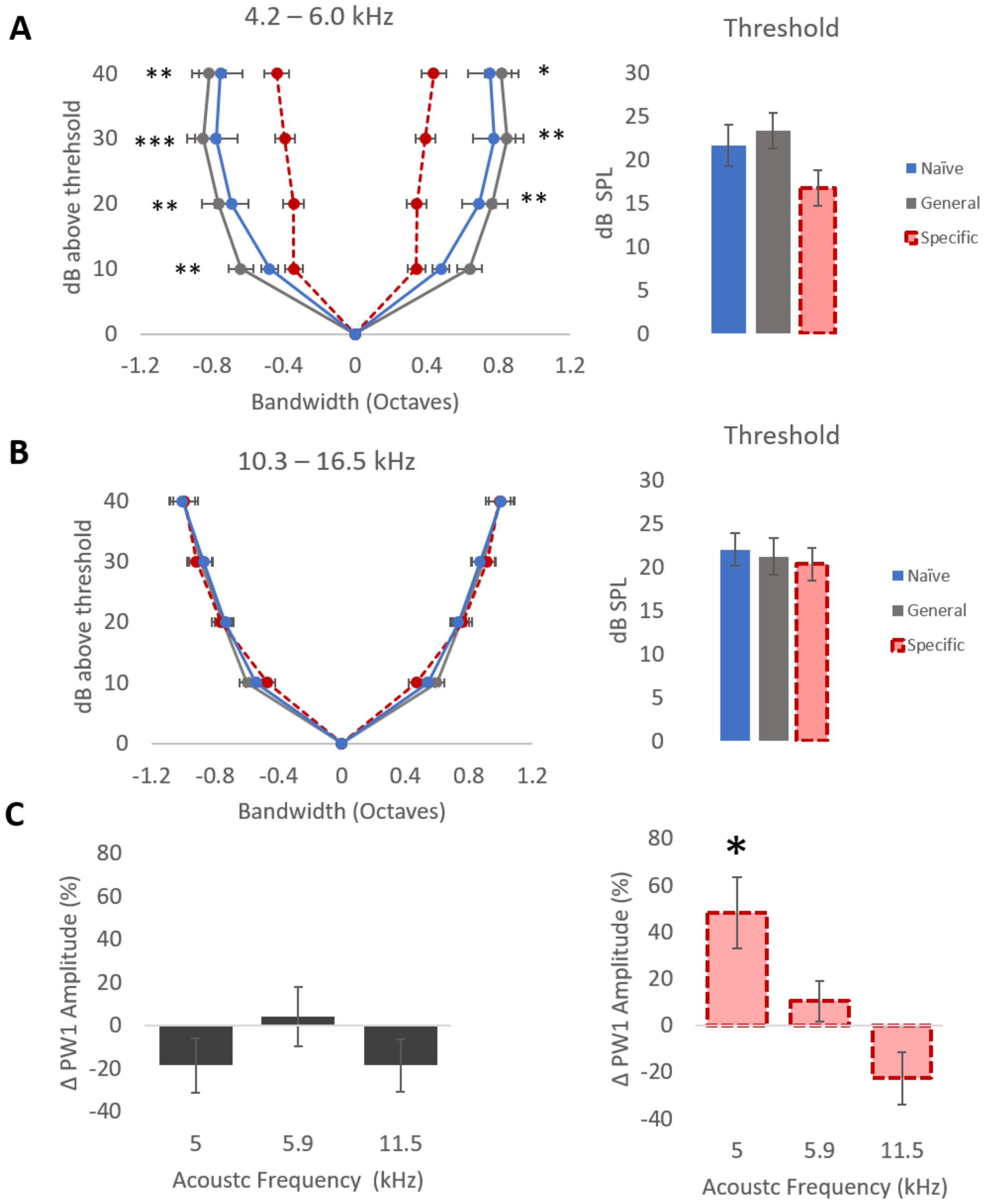


**Fig. S3.** *Frequency-specific memory is associated with signal-specific auditory system plasticity.* Panels represent sound-evoked neural responses from the auditory cortical (A,B) and auditory brainstem response (C) recordings. (A) For auditory cortical recording sites tuned near (within either +/- 1/3 octave) of the signal tone, animals with frequency-specific memory had significantly narrower tuning bandwidth animals with general memory and naïve animals. There were no differences between animals with general memory and naïve animals. There were no differences in cortical response threshold in any group. (B) For auditory cortical recording sites tuned far (1.03-1.33 octaves) away from the signal tone, there were no group differences in either tuning bandwidth or response threshold. (C) Analysis of learning-induced changes in PW1 amplitude revealed signal-tone specific amplitude increase in animals with frequency-specific memory with no significant amplitude changes in animals with frequency-general memories. *$p<0.05$ **$p<0.01$ ***$p<0.001$. In (A,B), asterisks on the left represent comparisons between general and specific memory; on the right, naïve vs . specific memory. No significant differences were found between naïve and general memory groups. All error bars represent +/-SEM.

**Tables:**

**Table S1.** *Summary of subgroup correlations*[1]

| | All | General | Specific | Vehicle | RGFP966 |
|---|---|---|---|---|---|
| 5.0 kHz: PW1 baseline amplitude *vs.* % Bar presses | 0.137 | 0.534 | 0.320 | 0.327 | 0.114 |
| 5.0 kHz: Δ PW1 amplitude *vs.* % Bar presses | 0.890*** | 0.485 | 0.934* | 0.766* | 0.975* |
| 5.0 kHz: % Bar presses vs. Cortical bandwidth | -0.668* | 0.738** | -0.594 | -0.454 | -0.740 |
| 5.0 kHz: Δ PW1 amplitude *vs.* Cortical bandwidth | -0.838** | -0.533 | -0.737 | -0.830* | -0.865 |
| 5.0 - 5.9 kHz: ΔΔ PW1 amplitude *vs.* Δ% Bar presses | 0.727* | 0.082 | 0.729 | 0.346 | 0.885 |
| 5.0 – 11.5 kHz: ΔΔ PW1 amplitude *vs.* Δ% Bar presses | 0.696* | 0.208 | 0.959** | 0.630 | 0.889 |

[1] This tables displays the Pearson r values for correlative data for the following groups of individuals: (1) all subjects, (2) subjects with frequency-general memory, (3) subjects with frequency-specific memory, (4) vehicle-treated subjects, and (5) RGFP966-treated subjects. *p<0.05; **p<0.01; ***p<0.001; ΔΔ: difference in the change in; Δ%: change in percentage
